# Exploiting cryo-EM structures of actomyosin-5a to reveal the physical properties of its lever

**DOI:** 10.1101/2023.03.19.533260

**Authors:** Molly S.C. Gravett, David P. Klebl, Oliver G. Harlen, Daniel J. Read, Stephen P. Muench, Sarah A. Harris, Michelle Peckham

**Affiliations:** Astbury Centre for Structural Molecular Biology, University of Leeds, Leeds, UK; School of Molecular and Cellular Biology, University of Leeds, Leeds, UK; School of Physics and Astronomy, University of Leeds, Leeds, UK; School of Biomedical Sciences, University of Leeds, Leeds, UK; School of Mathematics, University of Leeds, Leeds, UK

**Keywords:** Myo5a, CryoEM, motor protein, actin filaments, cantilever bending stiffness

## Abstract

Myosin 5a (Myo5a) is a dimeric processive motor protein that transports cellular cargos along actin filaments. Its long lever is responsible for its large powerstroke, step size and load-bearing ability. Little is known about the levers’ structure and physical properties, and how they contribute to walking mechanics. Using cryo-electron microscopy and molecular dynamics simulations, we resolved the structure of monomeric Myo5a, comprising the motor domain and full-length lever, bound to F-actin. The range of its lever conformations revealed its physical properties, how stiffness varies along its length and predicts a large, 35 nm, working stroke. Thus, the newly released trail head in a dimeric Myo5a would only need to perform a small diffusive search for its new binding site on actin, and stress would only be generated across the dimer once phosphate is released from the lead head, revealing new insight into the walking behaviour of Myo5a.

## Introduction

Myosins comprise a large family of cytoskeletal motors with a diverse range of biological functions ^1-3^. Each myosin comprises an N-terminal motor domain, which contains a nucleotide and an F-actin (filamentous-actin) binding site, followed by a lever, which comprises the converter and a light chain binding region, and the C-terminal tail comprised of specific domains that dictate the cellular function of the myosin. Small-scale conformational changes in the motor, powered by ATP, are amplified into a large-scale lever movement, which drives the movement of myosin along F-actin. Thus, myosin efficiently uses ATP hydrolysis to drive directional force and motion along filamentous actin (F-actin).

The mechanics of motors operating at the nanoscale are largely different to macroscopic motors, due to the viscosity of their environment, and the fact they are subject to thermal fluctuations and Brownian motion ^4,5^. There is only a limited understanding of how small-scale changes at the active site of the motor are translated into large scale coordinated movements of the lever. It is unclear what the range of lever conformations available to the motor is, how lever compliance varies along the lever and how these properties contribute to the lever movement and thus the working stroke. Here we address these questions using Myosin-5a (Myo5a), a widely expressed myosin ^6^ and provide evidence for the flexibility and conformational variability of the lever and how this contributes to its working stroke and walking behaviour.

Myo5a is a dimeric motor that walks processively along F-actin towards its barbed end ^7^, taking multiple steps without dissociating ^8,9^. Its two heads are dimerised by a coiled-coil tail, and it is recruited to cargo via its C-terminal cargo binding domain ^6^. Its long lever, comprised of the converter followed by an α-helix formed of 6 IQ (isoleucine-glutamine) motifs, each occupied by a calmodulin light chain (CaM), enables this myosin to take 36nm steps along F-actin ^10^. This step size matches the spacing of the F-actin helical pseudo-repeat and is thought to minimise azimuthal distortion ^8,9^. Its ability to take multiple steps (processivity) depends on its high duty ratio ^8,11,12^.

Although the step size of Myo5a is 36 nm, its working stroke was estimated to be only 25 nm ^12^, comprised of 20 nm from P_i_ release, followed by ∼5 nm from ADP release, which is the rate limiting step of its ATPase cycle (reviewed in ^7^). This discrepancy led to the idea that there needs to be an internal strain (or deformation) between the two heads to allow both heads to bind to actin at the same time, and that ADP release is a strain dependent step ^12^. Subsequently, it was shown that ADP release from the lead head of the dimer, bound to F-actin, is ∼250 times slower than that of the trail head ^13^. This biases ATP binding to the trail head followed by its detachment and search for the next binding site. Once this happens, the newly released head needs to perform a rapid diffusive search for its next binding site (estimated as ∼11nm in 100μs ^12^) before it can rebind.

The mechanical properties of the Myo5a lever are thus critical for its walking behaviour. It must be able to accommodate internal strain (or deformation) to co-ordinate and promote forward stepping, and able to resist a stall force of ∼3pN that prevents forward motion ^8^. To gain a better understanding of the mechanical properties of the lever, we have used cryo-EM to generate high resolution 3D structures of a single-headed Myo5a construct, comprised of the motor domain and a full length 6IQ lever (S1), bound to F-actin. By resolving multiple conformations of Myo5a S1 from a single cryo-EM dataset we have uncovered the intrinsic flexibility of the molecule, determined the lever stiffness in 3D, and identified regions of compliancy along its length. This allows us to interpret how lever flexibility influences the working stroke of Myo5a.

## Results and Discussion

### 3D cryo-EM classification captures 9 classes with levers in variable positions

To investigate the flexibility of the lever of Myo5a S1, we used cryo-EM to determine its structure when bound to F-actin in rigor. Previous cryo-EM studies of Myo5a-1IQ bound to actin ^14^ have shown a continuous conformational heterogeneity that was lower in rigor than in the ADP state. Thus, we chose rigor to optimize the chances of resolving multiple conformations of the lever. Focusing on a single Myo5a S1 and a single F-actin subunit allowed us to generate a 3D reconstruction that resolved the entire lever.

3D classification of Myo5a S1 molecules bound to F-actin resulted in 9 classes, in which density along the length of the lever could be observed (Fig. 1A). The position of the motor is unchanged between the classes while the position and conformation of lever changes from *a to i* (Fig. 1A). These classes likely represent snapshots of the lever across the range of continuous motion that the lever can undergo, reflecting the flexibility of the lever. Classes in which the lever angles are shallow are not due to compression in the ice as views contributing to the reconstruction were well distributed. The number of particles in each class is similar, and the global resolution of each class varied from 7.5 Å-10.7 Å (Fig. 1A).

**Fig. 1:**
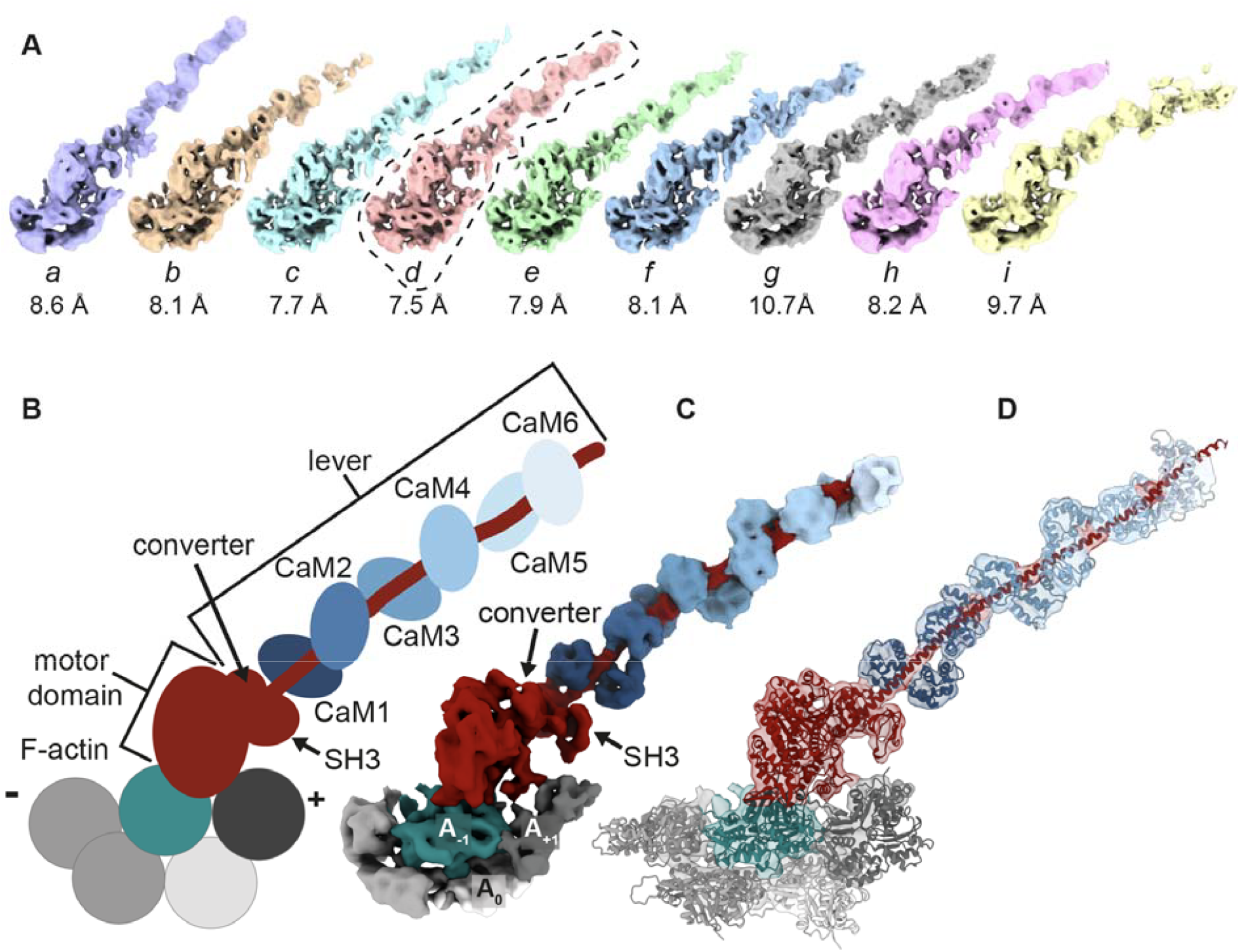
Cryo-EM shows the lever variability in Myo5a S1. **A:** Cryo-EM maps for the nine 3D classes generated following our analysis (contour level: 0.28). Classes are displayed in order of lever bend along the F-actin longitudinal axis. The estimate of the masked global resolution at 0.143 FSC (Fourier Shell Correlation) is displayed beneath each class. The circled class (*d*) is the class with the best resolution that was selected for generating the pseudoatomic model. **B:** Diagram showing the color scheme for F-actin, myosin heavy chain (red) and each of the 6 calmodulin light chains (CaMs, shades of blue) used in all figures. CaMs are labelled CaM 1-6, where the number refers to which IQ motif the CaM is bound. The polarity of F-actin is indicated by + for the plus and – for the minus end of the filament. The two adjacent F-actin subunits that interact with the motor domain are coloured green for the F-actin subunit closest to the minus end (A_-1_), and dark grey for the F-actin subunit closest to the plus end (A_+1_). **C:** Cryo-EM split map (contour level: 0.25) of a single Myo5a S1 class (class *d*) with the full-length lever (global resolution of 7.5 Å at 0.143 Fourier Shell Correlation (FSC)). **D**: The pseudoatomic model of Myo5a S1 is shown fitted into the split map (from **C)**.

Within each of the 9 classes, the resolution of the cryo-EM density map along the lever decreased moving away from the motor. For example, in class *d* the global resolution decreased from ∼7 to 25 Å, from the converter to IQ6 (Fig. S1B). This reflects conformational heterogeneity even within a single class, again demonstrating the flexibility within the lever. Using the cryo-EM data to perform an initial refinement that focused on a single motor domain and the first IQ produced a 4.2 Å reconstruction that was similar in structure to that previously reported for chicken Myo5a ^14^ with a single IQ motif bound to F-actin in rigor (Fig. S2). Small differences in the F-actin structure in our reconstruction compared to that in the earlier report ^14^ likely arise from our use of F-actin that was not phalloidin stabilized. This shows that the interaction of the motor with F-actin is relatively invariant.

The cryo-EM density for class *d* (Fig. 1C), which had the highest overall resolution, was fit with a pseudoatomic model (Fig. 1D) using all-atom molecular dynamics (MD) simulations, to determine the arrangement of the bound CaMs and their interactions within the lever. The conformation of the CaMs interacting with IQ motifs 1-5 appeared highly similar in the cryo-EM density. However, the N-lobe of CaM6 (CaM bound to IQ6) (Fig. 1C, 2D-E) is poorly resolved. It has previously been predicted that the CaM bound to IQ6 is in an extended conformation due to the substitution of Gly7 with Arg (Fig. 2F) ^15,16^. Thus, in the pseudoatomic model the extended light chain conformation of MLC1P bound to IQ4 in the yeast myosin-5 Myo2p crystal structure (PDB:1M46 ^17^) was used to build CaM bound to IQ6. The model additionally included the C-terminal FLAG tag sequence (DYKDDDDK), which is present after IQ6 (residue 908 onwards). The structure of the lever was then subjected to all-atom simulations in which the heavy chain was restrained, and the CaMs were unrestrained.

**Fig. 2:**
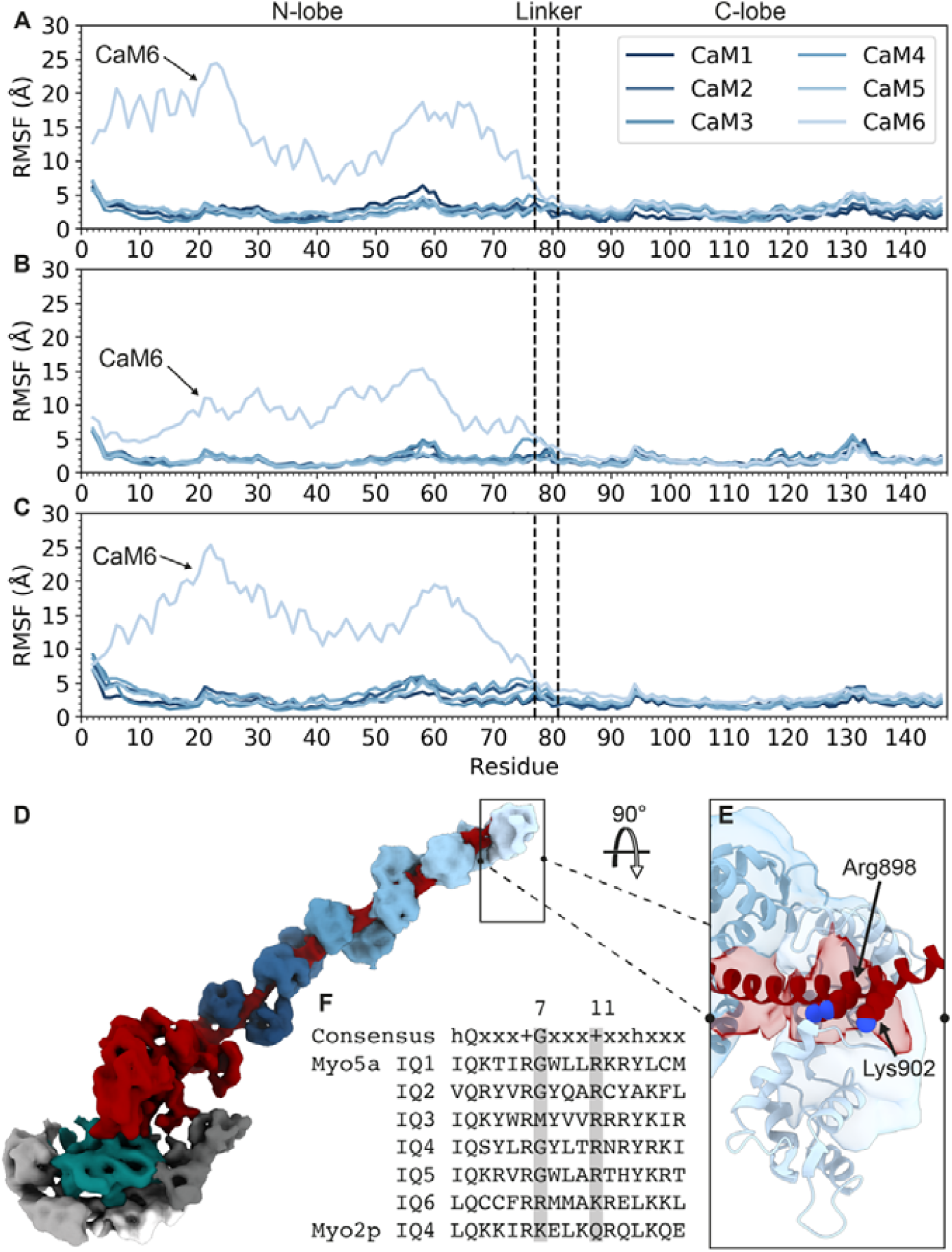
Variation in CaM interaction with IQ1-6 along the length of the lever. **A-C:** Plots of per residue RMSF (Root Mean Square Fluctuation) of CaMs obtained from 3 individual pseudoatomic model simulations in which the heavy chain is restrained and the CaMs are unrestrained. The locations of the N-lobe, linker and C-lobe are indicated. **A-C** show the results for these individual repeats. RMSF was calculated in CPPTRAJ (C++ Process Trajectory) ^42^. **D:** cryo-EM map of Myo5a S1 demonstrating the weak density for the N-lobe of CaM bound to IQ6 (CaM6) boxed. **E:** 90° rotation of boxed region in **D**, with a pseudo-atomic model fitted to the cryo-EM density. Residues at positions 7 (Arg898) and 11 (Lys902) of IQ6 are displayed in sphere mode and labelled. **F:** Sequence alignment of all 6 IQ motifs of murine Myo5a together with IQ4 of Myo2p (PDB:1M46 ^17^. Positions 7 and 11 of each IQ motif are highlighted in grey.

The substitution of Gly7 with Met in IQ3 of Myo5a (Fig. 2D & F) does not appear to result in an extended conformation in the bound CaM as density for both lobes can be seen. Interestingly, in a reference IQ motif, this substitution weakened its interaction with CaM by ∼10-fold ^15^. In the full[length intact lever, it is likely that the presence of neighbouring CaMs reduces the flexibility of the N-lobe and stabilizes its interactions with the IQ motif, suggesting CaMs bind the heavy chain cooperatively. It is possible that under some conditions this lobe could become extended, as a potential strategy for regulating myosin activity, by weakening the lever.

To confirm the extended conformation of CaM6 does indeed increase the mobility of the N-lobe, and that it does not tend towards the canonical closed state, we compared the mobility of CaM6 in the MD simulations to the other 5 CaMs (Fig. 2A-C). The mobility of the N-lobe of CaM6 was much higher than that for CaMs 1-5, consistent with its weak density in the cryo-EM map (Fig 2D), while that for the C-lobe was similar. Substitution of the conserved Gly residue at position 7 in the IQ motif to a bulkier Arg residue in IQ6 (Fig. 2D-F) could account for this weaker interaction, as previously observed in a reference IQ motif where this weakened its interaction with CaM by ∼3-fold ^15^ and for the Gly7 to Lys substitution in IQ4 of Myo2p ^17^. The recent shutdown structure of full length Myo5a shows all 6 IQ-CaM interactions are conserved ^18^, however, the N-lobe of CaM bound to IQ6 in each lever is in close contact with one of α-helical strands of the coiled coil, stabilizing the sharp bend at the head-tail junction ^18^. It is possible that in Myo5a S1, the lack of downstream sequence is partly responsible for weaker interactions between the N-lobe of CaM and IQ6 compared to the full-length motor. Overall, it is likely that the N-lobe of CaM bound to IQ6 is labile, which may be important for aiding switching between the shutdown and active state, and that CaMs bound to IQ3 and IQ6 can extend into the solvent, as reported for Mlc1p and Myo2p at IQ4, when there is no downstream sequence.

### The stiffness of the Myo5a lever varies along its length

The decrease in local resolution along the lever and the 9 different lever positions in the 3D classes, demonstrate the intrinsic flexibility of the unconfined lever. Using the 3D classes, we calculated the overall bending stiffness of the lever, by assuming it to be a cantilever beam held at one end by the stable interaction of the motor with F-actin, using beam theory from engineering (Fig. 3, Fig. S3). This approach measures the difference in position of the distal end of the lever in each 3D cryo-EM class from the mean position (Fig. S3A) (see Materials and Methods for details). We define the start of our lever as the converter and the end as the midpoint along the CaM bound to IQ6 (Fig. S5A). As there is only density for the C-lobe of CaM6 we used the conformation of CaM2 in IQ1-2 crystal structure (PDB: 2IX7 ^19^) to model the N-lobe and keep all CaMs equivalent.

**Fig. 3:**
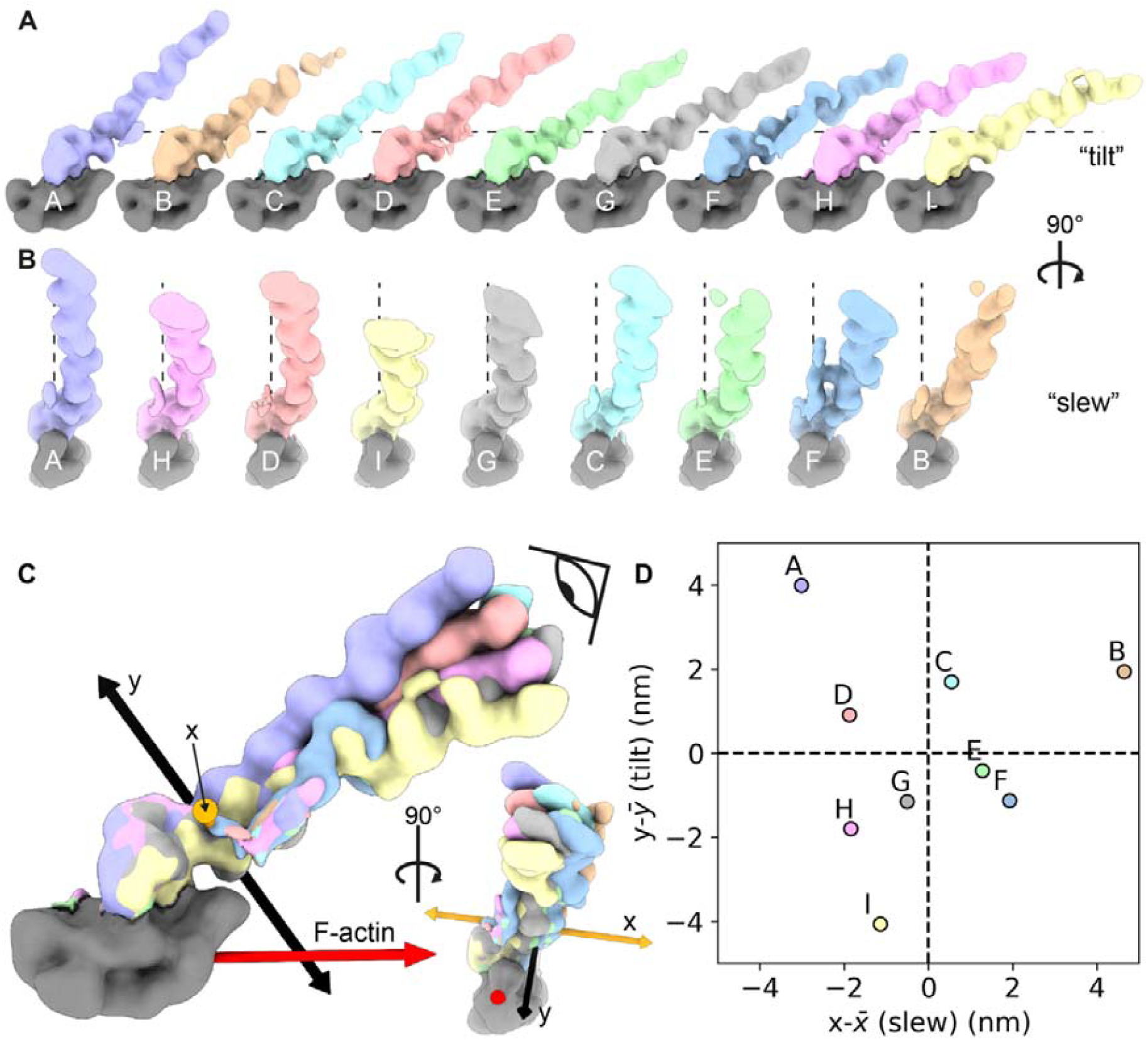
Lever flexibility is directionally isotropic within cryo-EM 3D classes. **A:** Post-processed maps of cryo-EM 3D classes in order of lever bend along the F-actin longitudinal axis (tilt). Visualized in the xy plane of the maps. The dashed line parallel to the x axis of the maps emphasizes the change lever bend between classes. Maps are Gaussian smoothed (SD 5 Å) (contour level: 0.15). **B:** Post-processed maps of cryo-EM 3D classes in order of lever bend along the F-actin short axis (slew). Visualized in the zy plane of the maps. The dashed line parallel to the y axis of the maps emphasizes the change lever bend between classes. **C:** Post-processed maps of cryo-EM 3D classes overlayed. The coordinate system the cantilever bending stiffness measurements were taken from are displayed as 3D arrows (see Materials and Methods). x-axis = orange arrows, y-axis = black arrows, z = axis mean lever vector (see Materials and Methods). Eye shows viewpoint graphed in **D. D:** Plot of the displacement of the end of the lever in each class from the mean (z) (see Materials and Methods), used to calculate the cantilever bending stiffness. Changes in x represent motion across the F-actin short axis (slew), changes in y represent tilt motions towards and away from the F-actin longitudinal axis (tilt).

We calculated the cantilever bending stiffness to be 0.75 ± 0.07 pN nm^-1^ for Myo5a, and to be directionally isotropic, as the calculated values for slew (across F-actin short axis) and tilt (bending towards and away from F-actin) are not significantly different (0.77 pN nm^-1^ and 0.73 pN nm^-1^ respectively (± 0.3 pN nm^-1^ confidence interval)) (Fig. 3, Fig. S3)) and the covariance is small compared to the variance (Pearson’s correlation coefficient is 0.02). The calculated bending stiffness value is similar in magnitude to previous estimates from optical trap measurements (0.2 pN nm^-1 12^) and negative stain EM (ns-EM) images of dimeric actomyosin-5a (0.26 pN nm^-120^). However, those estimates only indirectly determine stiffness based on changes in F-actin, whereas this is the first analysis performed directly on the lever.

The cantilever bending stiffness allows the calculation of the maximum deflection of the lever when forces are applied to its end. It has previously been reported that the intramolecular strain across the dimer when both heads are bound is 2 pN, and that the stall force when applying a backwards load to the dimer is 3 pN ^8,21^. Additionally, when a 4 pN backwards load is applied to Myo5a S1 there is equal distribution between reversal of the powerstoke (pre-powerstroke position) and the post-powerstroke conformation ^21^. Based upon our bending stiffness value (0.75 pN/nm), a 2 pN load would deflect the lever by 1.5 nm, which corresponds to 0.9 nm along F-actin. A 4 pN load would deflect the lever by 3.0 nm, which is 1.7 nm along F-actin (see Materials and Methods). Interestingly, a 4 pN force would deflect the lever close to the limit of the unstrained fluctuations we observe (∼4 nm) (Fig. 3D). The lever deflections in this force range are approximately equivalent to the change in lever end distance between the rigor and ADP state. This could explain how forces in this range alter lever position sufficiently to prevent entry into the rigor state. Note that deflections close to motor will be much smaller, 0.04-0.09 nm for CaM1 (0.03-0.05 along F-actin).

The density for all 6 CaMs in our 3D classes further allows us to determine if the lever behaves as a beam with uniform bending or if there are distinct points of flexion along its length. These are thought to arise from increased spacing and reduction in CaM-CaM interactions between a CaM bound to an IQ motif that is 25 residues long and its neighbouring C-terminal IQ motif, 23 residues long, such as IQ2 and IQ3 (12). A further pliant region could also lie between the converter and the CaM bound to IQ1 (CaM1) (13, 14). If the lever behaves as a uniform beam, then the cantilever bending stiffness (0.75 pN nm^-1^) is given by 3*EI*/*L*^3^, where *EI* is the local bending stiffness and *L* is the average length between the midpoint of CaM bound to IQ6 and the converter in our data (20.4 nm). This corresponds to a local bending stiffness (*EI*) of 2100 pN nm^2^ if the bending stiffness is uniform along the length of the lever (dotted red line, Fig 4B). To quantify the local bending stiffness at points along the lever we first discretized Myo5a into 7 subdomains (Fig. 4A). We then used the variance in subdomain conformations captured in the cryo-EM 3D classes (Fig. S4A-C) to determine the local bending stiffness of these subdomains and compared those values to the value for a beam with uniform bending along its length (Fig 4B, S5F, and Table S1).

**Fig. 4:**
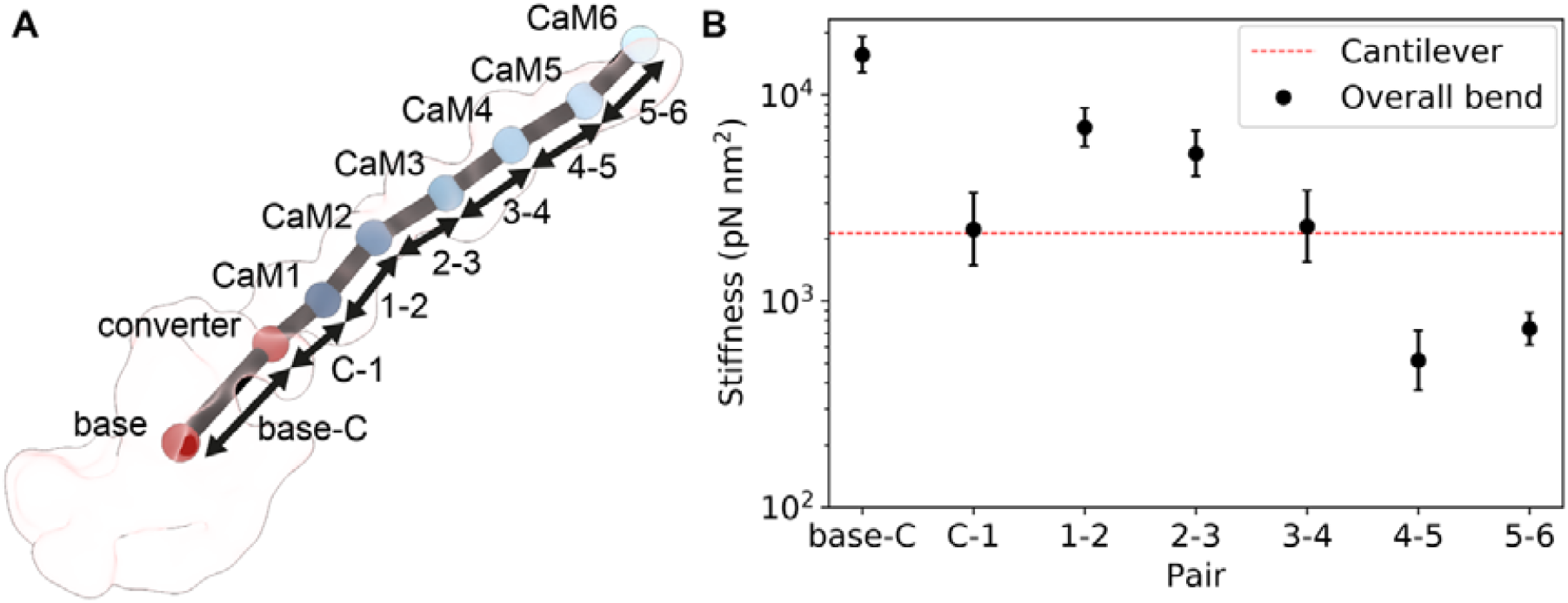
Determining the local bending stiffness of lever subdomains from the cryo-EM data. **A:** An example discretization of Myo5a S1 (class *d*) into 7 subdomains for calculating local bending stiffnesses (see Materials and Methods for details), displayed within the gaussian smoothed cryo-EM map (SD 5 Å) (contour level: 0.15). **B:** The calculated local bending stiffnesses for subdomains of Myo5a S1 (**A**). The expected local bending stiffness for a beam with uniform bending along its length calculated from the cantilever bending stiffness is plotted as a red line (Cantilever). Base = F-actin binding interface, C = converter, 1 = CaM1, 2 = CaM2, 3 = CaM4, 5 = CaM5, 6 = CaM6. Error bars show the SD of the random error (see Materials and Methods for details) as a percentage of the reported stiffness.

The local bending stiffnesses calculated from our cryo-EM data (Fig. 4B, Table S1) suggest that the lever stiffness is not uniform but varies along the length of the lever. As anticipated, one of the regions of increased pliancy is the ‘pliant point’ between the converter and CaM1 ^22,23^, which has a bending stiffness similar to that expected for a uniform beam. Thereafter, the regions between each IQ range in stiffness from stiffer than a uniform beam (IQ1-2 and 2-3), to equally as stiff (IQ3-4) and less stiff (IQ 4-5 and 5-6) (Fig. 4B). Our data suggests that the variation in IQ spacings and CaM-CaM interactions do not dictate the stiffness at subdomains in contrast to earlier predictions ^16^. Moreover, bending stiffness (Fig. 4B, Table S1, Fig. S5F) is not correlated with subdomain pair conformation as measured by angular distribution (Fig. S4A-C), distance (Fig. S4D) or interdomain interactions. Of note, measurements from previously obtained structures fall within a similar range to our measurements (Fig. S4). Finally, if Myo5a is a cantilever, the moment (tendency of a force to cause bending about the hinge) at each point along the lever is proportional to its distance from where the force is being applied. Thus, if there are indeed regions of pliancy along the length of the lever then the effects of forces on these regions will be most strongly felt the further away from the cantilever end and the closer to the opposing force of the fixed end they are. This may explain why only the region between the converter and CaM1 has been the previously identified as pliant, as previous EM analysis was performed on the double head bound dimer as opposed to an unrestrained lever considered here.

The change in local bending stiffness along the length of the lever is likely to be important for Myo5a mechanics. The lever must be sufficiently stiff to generate intramolecular strain and to withstand load from cargos without collapsing. The lever must also be sufficiently flexible to accommodate changes in step size, stereospecific binding to actin, and swapping filament track in a F-actin network ^20,24^. Myo5a constructs that comprise 2 IQ domains followed by a single alpha helix (SAH), which replaces IQ domains 3-6, can still walk processively, but do not generate intramolecular strain ^25^. The stiffness of a SAH is ∼150 pN nm^2^ which is ∼1/4 the stiffness between CaM4-6 (520 ± 200 pN nm^2^ at the CaM4-5 subdomain, and 730 ± 150 pN nm^2^ at the CaM5-6 subdomain). Taken together this could mean that: some rigidity at the start of the lever is required for transmitting the powerstroke and some rigidity (at least 4 x SAH stiffness) is required to communicate intramolecular strain between heads. The flexibility towards the end of the lever may help accommodate for stepping errors and angular disorder in F-actin ^20^.

### Lever flexibility augments Myo5a working stroke

Uncovering the range of motion of the lever enabled us to calculate the working stroke of Myo5a to be 35 ± 6 nm (mean ± standard deviation (SD)). To determine a population of lever end positions for the rigor state we fit the lever model of class *d* into each of the 3d classes (see Materials and Methods). Models of the ADP-P_i_ and ADP states were generated by superimposing the model of the lever for each 3D class (*a-i*, Fig. 1A) onto the converter domains of previously published Myo5a structures bound to an ADP-P_i_ analogue (ADP-VO_4_) (PDB:4ZG4, ^26^), and ADP (PDB: 7PM5, ^14^). As we lack a structure for the initial binding of Myo5a to F-actin in its ADP-P_i_ state, we modelled this state by docking residues 490-530 of 4ZG4 onto residues 492-532 of 7PM6, as the L50 domain is thought to bind actin first ^27^. We chose the end of the lever to be residue 912 (res 912 Cα atom), the last residue that is likely to interact with CaM in the active dimer. This residue was selected as it is the last residue the CaM bound to IQ2 interacts with in the IQ1-2 crystal structure (PDB: 2IX7 ^19^), and we do not have density for the N-lobe of CaM6 but believe it interacts with the heavy chain in the full-length dimer. We then determined the average distance travelled along the F-actin longitudinal axis between pairs of lever positions for every class superimposed onto the ADP-P_i_ model and onto the ADP model. This shows that the end of the lever translates 32 ± 5 nm (mean ± SD) along the F-actin longitudinal axis on average from the ADP-P_i_ to the ADP state. Extending this calculation to our rigor dataset, showed an additional translation of 3 nm along the F-actin longitudinal axis, between the ADP and the rigor state. This estimate of the working stroke is higher than that estimated previously (27 nm) using a similar approach ^26^, likely because we used a population of flexible levers rather than a single straight lever. Our estimate is also larger than that estimates obtained from optical trap measurements of the working stroke for dimeric Myo5a (25 nm) and S1 (21 nm) ^12^. These may be underestimates due to experimental uncertainties, such as where the myosin binds the nitrocellulose bead and how much applying force to the protein constrains the free motion of the lever.

Our larger estimate of the working stroke suggests that the diffusive search a newly detached myosin head needs to perform to rebind to F-actin at the next binding site in the walking dimer, is only ∼1 nm and not ∼10 nm as previously suggested ^12^. Instead, the length of the working stroke of a single head is sufficient to translate the molecule along F-actin the length of a pseudorepeat. This suggests that strain across the dimeric molecule is not generated during binding of the new lead head, but only after P_i_ release, as the lead lever is prevented from entering the post-powerstroke conformation by the trail head.

## Conclusion

3D reconstruction of the full-length Myo5a lever domain in 9 different conformations has revealed properties of the lever that contribute to its mechanics. Analysing the cantilever bending stiffness demonstrates that the stiffness of the lever is directionally isotropic. This means that the lever can accommodate intramolecular strain in a walking dimeric motor in all directions as it navigates a complex actin network. Analysing the local bending stiffnesses along the length of the lever, has shown its stiffness is not uniform but varies along its length, with the first pliant region between the converter and CaM1 and further pliant regions beyond IQ3. The initial region of the lever, comprising IQs 1-3 and their associated CaMs, is relatively stiff compared to the end of the lever. Interestingly, no single characteristic, i.e. the conformation of the subdomain or the number of interdomain interactions seems to correlate with bending stiffness. This suggests that either the combined influence of these features encodes the local stiffness at subdomains or features yet unexplored.

The variable flexibility along the length of the lever demonstrated here challenges previous hypotheses that assume the lever is a beam with uniform bending along its length. It is possible that 2D ns-EM images of the dimer with both heads bound to F-actin under strain have led to this idea, as they show a taut lever as opposed to one that can flex and bend (e.g. a tightrope is rigid under strain but is made of flexible rope), which prevents different properties along the length of the lever from being revealed. This highlights the value of using a single head with an unconstrained lever. Additionally, it is hard to determine whether, and by how much, the particles are distorted by both the stain and the carbon support film in ns-EM, which is significantly limited in resolution compared to cryo-EM. Previous analyses of Myo5a lever stiffness from optical trap and 2D ns-EM data have calculated lever stiffness by probing a change in or along F-actin. Due to 3D resolution along the length of the lever we have been able to probe properties of the lever directly for the first time. 3D cryo-EM reconstruction is therefore an important approach in estimating bending stiffnesses and can provide a more direct analysis than other techniques.

By adding further complexity to the model of the working stroke, using the ensemble of lever conformations within the cryo-EM 3D classes, we demonstrate that Myo5a S1 is capable of a longer working stroke (∼35 nm) than previously described (28.5 and 21 nm) ^12,26^. Its working stroke is thus closer to the length of the F-actin helical pseudorepeat than previously thought, suggesting that strain between the two attached heads in the walking motor is only generated once the motor has bound to F-actin, and P_i_ has been released. The nature of this strain, and its effect on the structure of the lead and trail heads will require cryo-EM structures of the Myo5a dimer as it walks along actin, a key challenge for the future.

## Materials and Methods

### Sample preparation

A murine Myo5a S1 construct (residues 1-907 followed by a FLAG-tag) and CaM proteins were purified as described ^28,29^, and stored in liquid nitrogen (LN_2_). Both were kindly provided by Howard White. After thawing stored Myo5a S1 additional CaM was added in a 2:1 molar ratio to ensure all IQ motifs were fully occupied. Rabbit skeletal muscle G-actin was purified as described ^30^. G-actin was dialyzed into 0.2 mM CaCl_2_, 0.5 mM DTT, 0.2 mM ATP, and 2 mM Tris-HCl, at pH 8.0, and stored in LN_2_. After thawing, G-actin was polymerized on ice, by first exchanging Ca^2+^ for Mg^2+^ in exchange buffer (final solution concentrations: 1 mM EGTA, 0.27 mM MgCl_2_) followed by polymerization in polymerization buffer (final solution concentrations: 25 mM KCl, 1 mM MgCl_2_, 1 mM EGTA, 10 mM MOPS, pH 7.0) overnight on ice.

### Grid preparation and cryo-EM data acquisition

Quantifoil R2/2 carbon Cu 300 mesh grids (Agar Scientific, Stansted, UK) were glow discharged in an amylamine vapour at 20 mA for 30 s (GloQube, Quorum Technologies Ltd, Laughton, UK). Directly before application, F-actin was sheared by being repeatedly drawn up and ejected with a gel-loading pipette tip to shorten filaments, to increase the amount of F-actin observed occupying grid holes. 1 μL of sheared F-actin (0.5 μM) was applied to the grid and incubated for 2 mins. 3 μL of Myo5a S1 supplemented with CaM (3.3 μM + 6.7 μM, respectively) was added to the grid in the Vitrobot Mark IV (Thermo Fisher, Altrincham, UK), followed by a second incubation of 2 mins. Final concentrations of proteins were: 0.125 μM actin, 2.5 μM Myo5a S1, 5 μM CaM. All dilutions were done using100 mM KCl, 0.1 mM EGTA, 1 mM MgCl_2_, 10 mM MOPS, pH 7.0. Grids were then blotted with Whatman no. 42 Ashless filter paper (Agar Scientific, Stansted, UK) for 3 s at force -25, 8□°C and 80 % humidity, drained for 0.5 s and flash-frozen in liquid ethane. Data was recorded on the FEI Titan Krios I (Astbury Biostructure Laboratory, University of Leeds, Leeds, UK) equipped with a FEI Falcon III detector operating in linear mode (Table S3).

### Cryo-EM image processing

First, MotionCor2 ^31^ was used to correct for beam-induced motion, and the contrast transfer function (CTF) was estimated using Gctf (GPU accelerated CTF) ^31^, before subsequent processing steps (Fig. S6). The start-end coordinates of F-actin filaments were manually picked using RELION 3.1 (Fig. S6B) ^32^. Particles were extracted in RELION 3.1 using helical parameters (box size 608 px, helical rise 27.5 Å, tube diameter 250 Å). The data was initially binned to 2.13 Å. Helical 3D refinement was used to produce an initial model (tube outer diameter 140 Å, angular search range tilt 15° and psi 10°, initial twist -166.15°, helical rise 27.5 Å, twist search -165° to -167° with a 0.1° step, rise search 26.5 Å to 28.5 Å with a 0.1 Å step) (Fig. S6C). A known structure of the Myo5a motor (PDB: 7PLU, ^14^) was rigid fit into the helical refinement map at a motor domain that was the best resolved and positioned at the centre of the box, using Chimera ^33^. A map of the fitted PDB structure was generated in Chimera ^33^, and used as an input to generate a wide mask of the motor domain in RELION 3.1 for masked 3D classification ^32^. Masked 3D classification was performed in RELION 3.1 to classify out undecorated actin and to only include particles with myosin bound at the centre of the box (5 classes, no image alignment, regularization parameter T = 4) (Fig. S6D). This dataset was then un-binned (1.065 Å) as initial reconstructions were reaching the Nyquist limit. 3D helical refinement followed by masked post-processing of this subset of particles produced a map with a 3.8 Å global resolution, but with limited detail across the lever (Fig. S6C). All global resolutions were determined using the gold standard Fourier Shell Correlation (FSC) reported to FSC = 0.143 (FSC_0.143_) using RELION 3.1. To compare the motor domain to the previously published chicken actomyosin-5a rigor structure (PDB: 7PLU ^14^), particle subtraction was performed subtracting all density outside of a mask comprising of the motor, the first 2 CaMs and 3 F-actin subunits. 3D refinement followed by post-processing, produced a map with a global resolution of 4.2 Å according to the FSC_0.143_-criterion (Fig. S6), which was similar to that previously published (3.2 Å, PDB: 7PLU ^14^) (Fig. S6E). This map was then locally sharpened using DeepEMhancer ^34^.

To improve lever resolution, particle subtraction was required to aid particle alignment and centralize the lever domain within the box. It was noted from initial 3D helical reconstructions that the lever density was smeared, so a wide mask containing 1 actin subunit, the motor, and a cone shape for the lever to accommodate flexibility was used for subtraction (Fig. S6F). A map was generated in Chimera comprising a single actin subunit, a motor (both from PDB: 7PLU, ^14^), and multiple copies of a Myo5a lever model (PDB: 2DFS ^35^) arranged in a cone shape tapering at the motor and splaying towards the lever end. These were positioned so that the boundaries of the cone met the density of the neighbouring actin bound heads in the helical map. A wide cone-shaped mask was generated in RELION 3.1 using the cone-shaped map (Fig. S6F). The signal outside of this mask was subtracted from the binned (to 2.13 Å) 2D images, and particles were re-centred bringing the lever to the focal point of the box. 3D refinement of the subtracted particles produced a map (cone subtracted map) with 4.3 Å global resolution according to the FSC_0.143_-criterion (Fig. S6G). This map was locally sharpened using DeepEMhancer ^34^. The cone subtracted map had improved resolution across the lever, however defined density for CaMs 5 and 6 could not be seen (Fig. S6G).

To resolve CaMs 5 and 6, 3D classification of the lever domain using a cone shaped lever mask was performed. This mask was generated in the same way, but the actin subunit and motor domain were excluded to classify based on lever conformation only. Attempts to classify into 10 or 20 classes both resulted in ∼9 reasonable classes (Fig. S6H). These conformations reconstructed by 3D classification are not distinct conformational states but representative of continuous thermally driven motion of the lever, as the conformation of the classes produced differed in both modes of division (10 or 20). Much larger numbers of particles may have resulted in a larger number of classes. The maps were locally sharpened using DeepEMhancer ^34^. Though classification led to a reduction in overall resolution due to loss of particles in the reconstruction, for the first time we were able to see across the full length of the lever while the motor is actin bound (Fig. S6I). However, we cannot rule out that there may be particles representing more extreme lever conformations outside of the subtraction and classification masks used to focus the image processing.

The cryo-EM 3D class with the best global resolution (7.5 Å) (Fig. S1A) was selected to build a pseudoatomic lever model (Fig. S6I). The local resolution of this map was calculated using SPOC (statistical post-processing of cryo-EM maps) (Fig. S1B), as the local resolution calculations in RELION are unreliable at resolutions lower than 10 Å ^36^.

### Cryo-EM model building and refinement

An initial lever model including the converter + 6IQ motifs sequence (698-907) + 8 residue FLAG-tag, and 6 CaM sequences, was built in Alphafold 2.0 using Collabfold Google collab notebooks ^37^. This model was then flexibly fit into the density of the best 3D class (*d*) using Isolde ^38^, applying distance and torsional restraints taken from murine IQ1-2 and CaM1-CaM2 structure (PDB: 2IX7, ^19^) to each CaM pair. Distance and torsional restraints from the Myo2p 25-residue spaced pair structure (PDB: 1N2D,^16^) were applied to interacting residues in 25-residue spaced pairs (Glu14-Arg91, in CaM2-3 and CaM4-5), as density corresponding to these interactions could be seen in the selected class map. Only the C-lobe of CaM6 was included in fitting as only density for this half of the molecule could be seen in the map. To model the N-lobe of CaM6 a homology model of CaM in an extended state, based on a Myo2p-Mlc1p structure (PDB: 1M46, ^17^), was built using Swiss-model ^39^. The C-lobe of the homology model was superimposed onto the C-lobe of the initial CaM6 model. The N-lobe (residues 3-84) of the homology model was then joined to the C-lobe of the initial CaM6 model and minimized in Isolde without map weighting.

A pseudoatomic lever model was generated by performing all-atom MD simulations of the fitted lever model to gain side-chain conformations. All simulations were performed using Amber20 ^40^ with the FF19SB forcefield ^41^. The lever model was protonated according to the Amber residue templates and then solvated with TIP3P water molecules in an octahedral box that extended 14 Å from the protein. K^+^ ions were added to neutralize the system, then KCl was added to a final concentration of 100 mM. After initial energy minimization the system was heated to 300 K as positional restraints were decreased from 100 to 0 kcal/mol/Å^2^, except for across the lever heavy chain. A restraint of 1 kcal/mol/Å^2^ was applied to the backbone of the lever heavy chain throughout the minimization, equilibration, and production runs, to permit CaM motion and interaction with the heavy chain side-chains but maintain the lever position seen in the cryo-EM map. Minimization and equilibration steps were performed on the ARC4 standard nodes (Intel Xeon Gold 6138 CPUs (‘Sky Lake’)). NMR distance restraints were also applied between interactions visible in the cryo-EM density (Glu14-Arg90). To do this, distance restraints were applied between the Cδ atom of Glu, and Cζ of Arg, to not dictate which N and O interact. These were weighted at 20 kcal/mol/Å^2^ within 1.9 Å of the bounds of the flat well restraint (3.4-5.3 Å), and a 20 kcal/mol/Å^2^ harmonic potential was applied outside of this range. The MD production runs used the pmemd.cuda module from Amber20 and were run on Bede using Tesla V100 GPUs. MD was performed for 300 ns in triplicate repeat. The Berendsen Thermostat was used as recommended to maximize GPU performance. To compare the flexibility of each CaM, per residue root mean square fluctuation (RMSF) analysis was performed using CPPTRAJ on each CaM for all three repeat simulations ^42^.

Following simulation, the average conformation was calculated in VMD (Visual Molecular Dynamics) ^43^, and the frame of the trajectory with the lowest global RMSD (Root Mean Square Deviation) with the average conformation was selected as a model for each repeat. A further round of minimization was then performed on the ARC4 general nodes to restore side-chains to low energy conformations. After minimization, each model was scored in MolProbity ^44^, and the repeat with the lowest MolProbity score and the largest proportion within the cryo-EM density was selected as the final lever model.

To generate a model of the motor domain, a homology model of murine Myo5a was made using the chicken actomyosin-5a rigor structure (PDB:7PLU ^14^) as the template in Swiss-model ^39^. This was then fit into the cryo-EM density using Isolde ^38^, applying distance and torsional restraints based on the template. Residues 1-698 were then joined to the pseudoatomic lever model (699-915). The regularize zone tool in Coot was used to correct over the stitch region ^45^. The F-actin subunits from the chicken actomyosin-5a rigor structure (PDB:7PLU ^14^), with phalloidin removed, were also fit into the density corresponding to actin using Isolde ^38^. As the resolution of F-actin and the motor domain was insufficient to fit side-chains, and their structures have already been published at high-resolution, only the backbone is included for these domains in our model ^14^. Side-chain orientations are included for the lever from the simulated data. The quality of the final model was assessed using MolProbity ^44^ (Table S4).

### Flexibility analysis

All flexibility analysis was performed on discretized representations of Myo5a S1. First the Isolde generated lever model (pre-simulation) and the homology model of the motor were flexibly fit into each cryo-EM class with torsional and distance restraints applied using Isolde ^38^. A coarse-grained representation of the conformation of Myo5a S1 was then defined by discretizing into 8 vectors (Fig. 4A & Fig. S5A) drawn between Cα atoms for the base (actin binding interface) (res 384 and 543), the converter (res 721 and 760), and CaMs 1-6 (res 137 and 64) (Fig. S5A).

The cantilever bending stiffness of the entire lever was calculated using the variance in lever displacement from the mean. First, a vector representing the lever was drawn between the midpoint of the vector describing the converter, and the midpoint of the vector describing CaM6 for all classes (Fig. S5A). The mean of these lever vectors was then to define a new coordinate system with an origin at the midpoint of the vector describing the converter (Fig. 3C) and the *z*-axis along the mean direction of the lever. The *x* and *y* axes in the perpendicular plane were defined such that changes in *y* represented tilt motions (towards the F-actin longitudinal axis) and changes in *x* presented slew motions (perpendicular to the F-actin axis). Thus, the y-axis was orthogonal to *z*, and the *yz* plane was parallel to the vector representing F-actin (res 179 Cα atom of the 0^th^ and 13^th^ subunit of a model F-actin). The model of F-actin was made by fitting an actin subunit from a previously published structure into the 0^th^ and 13^th^ subunit of our initial F-actin helical reconstruction (Fig. S6C). Then for each class, the distance, *d*_c_, in the *xy* plane between the end point of the lever unit vector and the end point of the mean unit vector was calculated for each class (Fig. S3). The variance in lever conformation was calculated using Equation (**1**),

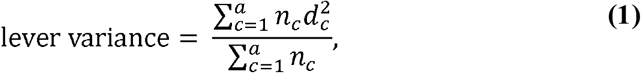

where *n*_c_ is the number of particles in the class. This was done for the overall displacement, and also for displacement in a particular direction (tilt and slew, along the y- and x-axis respectively), to calculate the overall bending stiffness (Fig. S3B) and the bending stiffness in both directions (tilt/slew, Fig. S3C).

The overall cantilever bending stiffness (*k*_ov_) was calculated using Equation (**2**),

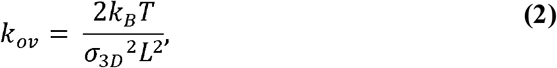

where *k*_B_ = 1.38 × 10^−23^ N m K^-1^ (Boltzmann constant), *T* = 281 K (temperature of grid making in Kelvin), *L* is the mean length of the lever vectors (20.4 nm), and σ3D^2^ is the variance calculated from the overall displacement within the 2 degrees of freedom. A factor of 2 is included in Equation (**2**) to account for the two degrees of freedom in bending along two perpendicular directions.

We also calculated a separate directional cantilever bending stiffness (*k*_*dir,i*_) for displacements in the *x* and *y* directions separately using Equation **(3)**,

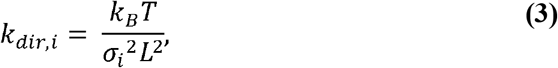

where 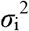 are the variances calculated from the displacements in *x* or *y*.

The Pearson’s correlation coefficient for a sample population was determined using Equation (4),

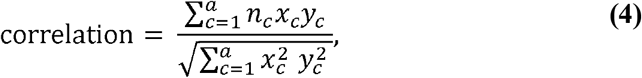

where (*x*_c_, *y*_*c*_) is the displacement between the class and mean unit vector in the *xy* plane.

To estimate the error in the calculation of the cantilever bending stiffness we calculated the effect of varying the value for displacement of the lever within reasonable bounds, given the grouping of molecules into classes. The values for displacement from the mean of each class were sorted into ascending order. A new value for this displacement was generated for each class, chosen randomly between the displacement values either side of it in the sorted list. The ranges for values at the start and end of the list were calculated using the value ± the distance from the single neighbour. The randomly generated displacements were used to calculate the unweighted variance using Equation **(5)**,

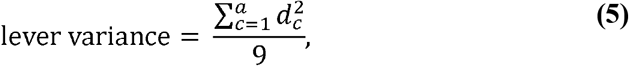

where *d*_c_ is the randomly generated displacement. This variance was used to calculate the cantilever bending stiffness using Equation **(2)** for the overall bending stiffness and Equation **(3)** for the directional bending stiffness in each direction. This process was repeated 1000 times, generating a new estimate of the cantilever bending stiffness each time. The SD of these randomly obtained bending stiffnesses was quoted as the error.

To determine the local conformation of lever subdomains in each class, the 8 vectors representing Myo5a S1 were paired into 7 subdomains. Local material axes *uvw* were defined for each subdomain (Fig. S5A-E), where the *u* axis is the vector between the midpoints of each vector in the subdomain vector pair and the *v* and *w* are in the orthogonal plane are defined such that the first vector of the subdomain pair lies halfway between *v* and *w* (*i*.*e*. at an angle of 45°). The angles between the subdomain vector pairs were then calculated in each plane (*vw, vu* and *wu*), θ_*vw* =_ θ, θ_*vu* =_ θ_1_-θ_2_, θ_*wu* =_ θ_3_-θ_4_ (Fig. S5C-E, Fig. S4A-C), the angles for θ and θ_1-4_ were taken from -90 to 90°. To determine if the distance between CaM interacting regions has an influence on local bending stiffness, the distance between the Cα atoms of known interacting residues in the N- and C-lobe of consecutive CaMs was calculated (Ser17 and Asn111, respectively) (Fig. S4D).

To calculate the local bending stiffnesses at subdomains (*k*_*sub*_), the variance (σ^2^) in angle between subdomain vector pairs within the classes (weighted by the number of particles in each class), was calculated in 3 orthogonal planes (*vw, vu, wu*) (Fig. S5A-E, Fig. S4A-C). The variation in the *vw* plane describes torsion (Fig. S5C), and the *vu* and *wu* planes describe bending stiffness in 2 directions (Fig. S5D-E). *k*_*sub*_ in each plane was calculated using Equation (6),

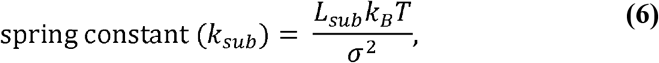

where *L*_*sub*_ is the average length of the subdomain in the cryo-EM 3D classes (the distance between the midpoints of each subdomain vector pair, Table S2) and σ2 is the variance in angle between subdomain vector pairs within the classes. The overall bending stiffness was calculated by averaging the values in the *vu* and *wu* plane.

To estimate the error for the local bending stiffness calculation at each subdomain we calculated the effect of randomly varying the angles between the subdomain vector pairs within reasonable bounds, given the grouping of molecules into classes, as follows. For each subdomain, the angles calculated between the vector pair in each class were sorted into ascending order. A new value for this angle was generated for each class, chosen randomly between the angle values either side of it in the sorted list. The ranges for values at the start and end of the list were calculated using the value ± the distance from the single neighbour. The variance (σ^2^) was calculated for these randomly assigned angles and used to calculate a bending stiffness using Equation (6). This process was repeated 1000 times, generating a new estimate of the local bending stiffness each time. The SD of these randomly obtained local bending stiffnesses was quoted as the error.

To determine the maximum deflection of the lever from its average position when a force is applied to the end, the cantilever beam deflection Equation (7) was used,

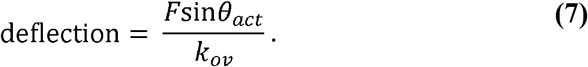

Here the applied torque is *F*sinθ_act_, where *F* is the force applied, and θ_*act*_ is the angle between the mean lever position and F-actin (Fig. S7A). As in optical trap studies the force is parallel to the F-actin axis, it was necessary to determine the deflection when an off-axis (not in the *xy* plane Fig. 3C) force was imposed using *F*sinθ_*rig*_.

The equivalent distance of the lever deflection along the F-actin axis was also determined using Equation (8),

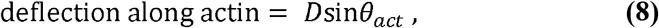

where D is the deflection in the *xy* plane Fig. 3C.

The deflection of the cantilever beam at a specific location was determined using the cantilever lever beam deflection Equation (9),

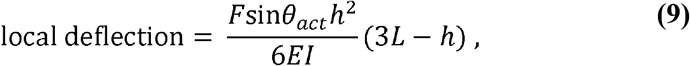

where *EI* is the local bending stiffness and *h* is the distance along the lever from the fixed end (the converter in our model).

Finally using cantilever beam theory, the moment at any point can be calculated with Equation (10),

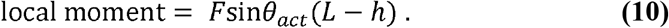

### Predicting the working stroke

The ensemble of lever conformations in the cryo-EM 3D classes were used to predict the working stroke of Myo5a S1. To determine the distance the working stroke translates the end of the lever along F-actin longitudinal axis, a vector between the 0^th^ and 13^th^ subunit (Cα, residue 179) of F-actin was calculated from an F-actin model (as above). The lever models generated from each cryo-EM class were superimposed onto the converter of a pre-powerstroke (PDB: 4ZG4, ^26^), and post-powerstroke ADP bound structure (PDB: 7PM6, ^14^). As 4ZG4 is not actin bound, to model the motor bound in the ADP-P_i_ (pre-powerstroke) state the L50 domain (residues 490-530) of 4ZG4 was superimposed with the actin interacting region of the L50 domain (residues 492-532) of 7PM6, as the L50 domain is thought to bind actin first ^27^. As we do not have density for the CaM6 N-lobe in our Myo5a-S1 map, but believe it interacts with the heavy chain in the full length motor, we use the equivalent residue in IQ6 (res 912) to the last residue in IQ2 that CaM interacts with in the CaM bound IQ1-2 crystal structure (PDB: 2IX7 ^19^) as our lever end. Vectors were drawn between the lever end of each class model of the ADP-P_i_ state to every other class model of the ADP state (Fig. 5A). The translation of the lever end along F-actin was calculated using Equation (11),

**Fig. 5:**
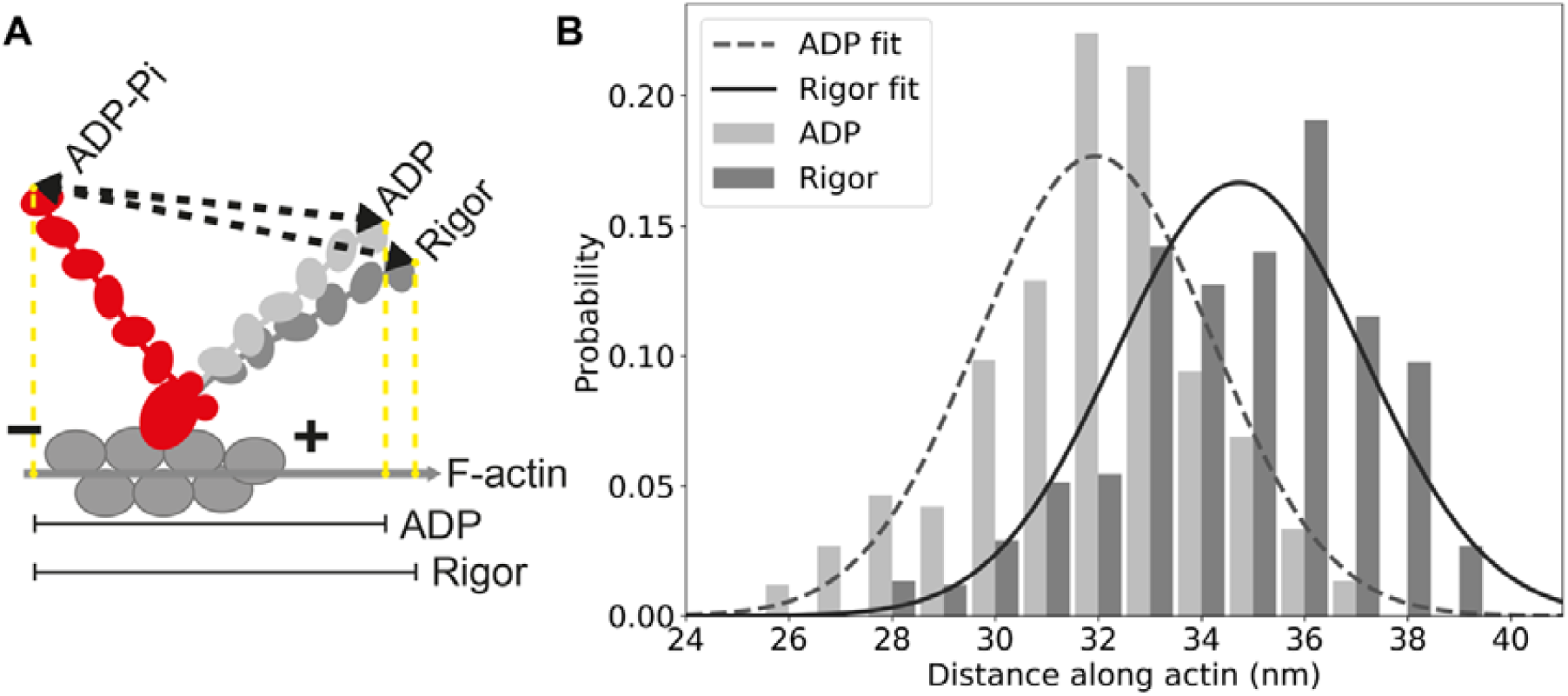
The working stroke of Myo5a is 35nm. **A:** Schematic diagram to demonstrate how distances for **B** were calculated. Distances between lever ends in the modelled ADP-Pi, ADP, and rigor conformations (see Materials and Methods for details) are shown as black dashed arrows. Yellow dashed lines show these distances as a translation along the F-actin vector (0^th^ to 13^th^ subunit, grey arrow, see Materials and Methods for details). + indicates the plus end of F-actin, -indicates the minus end of F-actin. **B:** Histogram of the translation of Myo5a lever ends along F-actin vector from the ADP-P_i_ conformation to ADP and to rigor (see Materials and Methods for details), and the fitted Gaussian distributions. The probability of each class being paired was calculated using the proportion of particles in each of the cryo-EM 3D classes (see Materials and Methods).

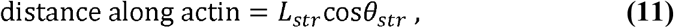

where *L*_*str*_ is the magnitude of an ADP-Pi to ADP lever end vector and θ_*str*_ is the angle between this vector and the F-actin axis (Fig. S7B). This was then repeated to calculate the working stroke from the ADP-P_i_ state models and our rigor models. The probability of a combination of classes being paired was calculated by taking the probability of an individual class occurring, using the fraction of particles in each class out of the total number of particles. The probabilities of each class in a pair were then multiplied to give the probability of that combination of classes being paired.

## Acknowledgments

We would like to thank Eva Forgacs, Betty Virok and Howard White for the generous gift that was the purified Myo5a S1 protein. We would like to thank Glenn Carrington, Peter J. Knight and Yasuharu Takagi for the helpful discussion and advice. We would like to thank the Astbury Biostructure Laboratory for their support. The FEI Titan Krios microscopes were funded by the University of Leeds (UoL ABSL award) and Wellcome Trust (108466/Z/15/Z). This work made use of the facilities of the N8 Centre of Excellence in Computationally Intensive Research (N8 CIR) provided and funded by the N8 research partnership and EPSRC (EP/T022167/1), through HECBioSim (EP/R013012). The center is coordinated by the Universities of Durham, Manchester and York, UK. This work was also undertaken on ARC4, part of the High-Performance Computing facilities at the University of Leeds, UK. Molly S. C. Gravett and David P. Klebl were PhD students on the Wellcome Trust 4-year PhD program (102174/B/13/Z) in The Astbury Centre funded by The University of Leeds. Michelle Peckham is funded by the Wellcome Trust Investigator award ‘Myosin structure and regulation’ (223125/Z/21/Z).

## Author contributions

Conceptualization: MSCG, SAH, SPM, MP,

Methodology: MSCG, DPK, OGH, DJR, SAH, SPM, MP

Investigation: MSCG, Supervision: SAH, SPM, MP

Writing—original draft: MSCG, MP, SPM, SAH

Writing—review & editing: MSCG, DPK, OGH, DJR, SAH, SPM, MP

## Data and materials availability

the EM density maps for 9 3D class reconstructions (Fig. 1A) have been deposited in EMDB: EMD-16846 (class *a*), EMD-16848 (class *b*), EMD-16849 (class *c*), EMD-16850 (class *d*), EMD-16851 (class *e*), EMD-16853 (class *f*), EMD-16852 (class *g*), EMD-16854 (class *h*), and EMD-16855 (class *i*). The simulated pseudoatomic model for class *d* (Fig. 1D) was deposited PDB 8OF8. The reconstruction focusing on the motor domain plus 2IQs (Fig. S6E) was deposited in the EMDB under EMD-16856. All data are available in the main text or the supplementary materials.

