## Supplemental Data for "Exploiting cryo-EM structures of actomyosin-5a to reveal the physical properties of its lever"

Molly S.C. Gravett *et al.*

**This PDF file includes:**

Figs. S1 to S7
Tables S1 to S4

**Figure S1: cryo-EM map resolution (class *d)***

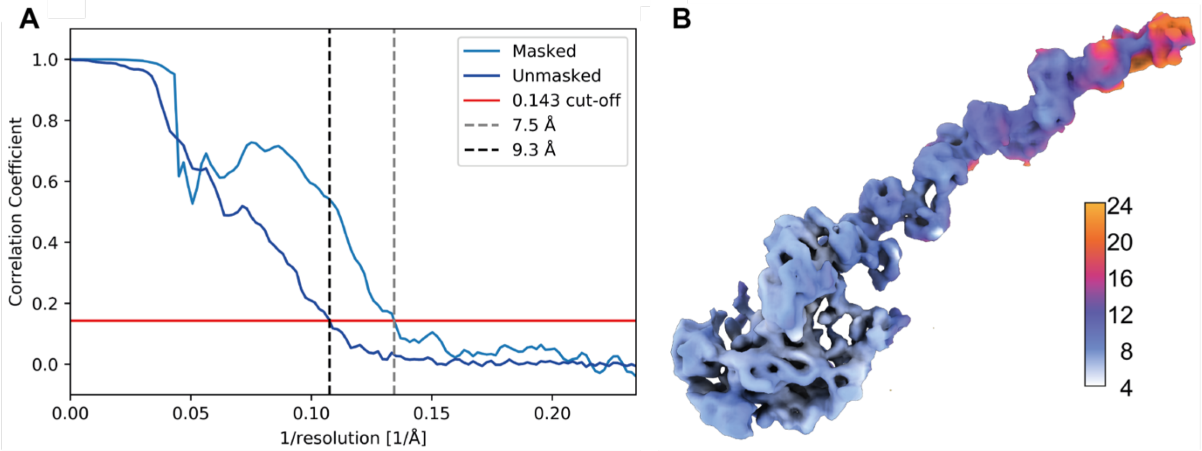


**A:** Fourier shell correlation (FSC) curve illustrating masked 7.5 Å resolution and unmasked 9.3 Å resolution at 0.143 FSC. **B:** Local resolution calculated with SPOC (Statistical Processing of Cryo-EM maps) [36] displayed on the F-actin bound S1 cryo-EM map (contour level: 0.25) (Fig. 1A.*d*). The color bar shows the resolution in Å.

**Figure S2: Fitting of a published rigor motor and CaM structure into the cryo-EM motor map obtained in this study**

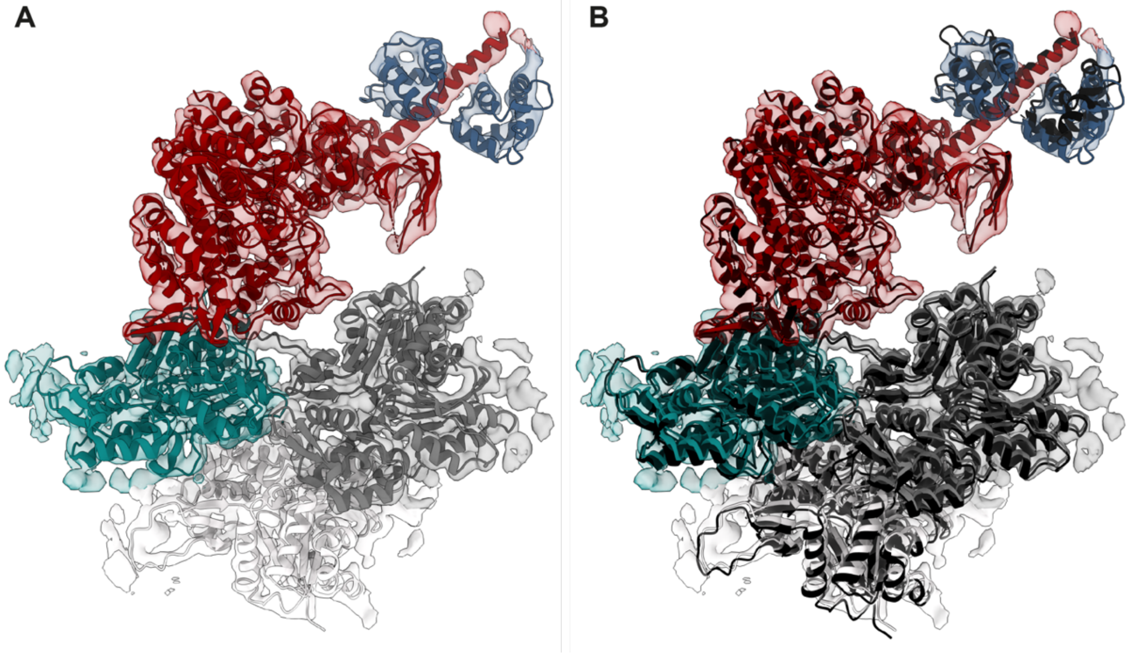


**A:** Cryo-EM map of murine Myo5a focusing on the motor + 2IQ and 3 F-actin subunits, contoured to show 1IQ. Individual chains of the previously solved chicken actomyosin-5a rigor model (PDB: 7PLU,[14]) were rigid fit into this cryo-EM density map. The essential light chain (as in 7PLU) was substituted with the CaM from the CaM-IQ1 crystal structure (PDB: 2IX7, [19] ). Colors as in Fig. 1B. **B:** As in **A**, but with the previously reported structure (7PLU,[14]) rigid fit into the density by the motor domain only (displayed in black). All rigid fitting was performed in ChimeraX [33]. Minor differences in the orientation of the motor relative to F-actin in our reconstruction can be seen by comparing the arrangement chains individually fit into our reconstruction with the original chicken actomyosin-5a structure. This is unlikely to be significant and could arise from sequence variation between chicken and murine Myo5a, and/or the use of phalloidin to stabilize F-actin in the earlier study [14].

**Figure S3: Calculating lever displacement for cantilever bending stiffness**

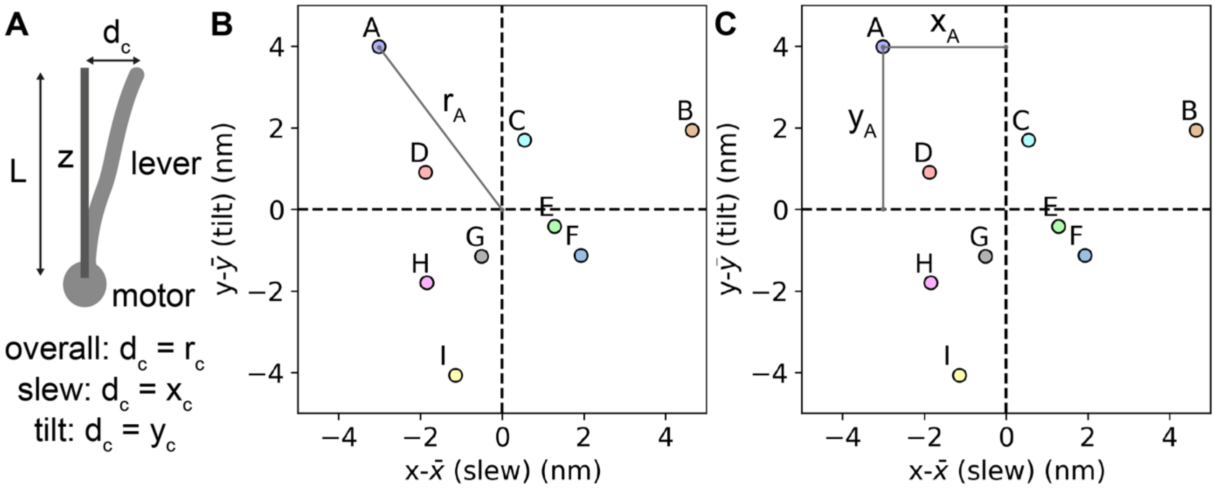


**A:** Schematic of cantilever-type bending of the lever of Myo5a. $d_c$ is the displacement of the lever in each class (c) from the mean position (z). L is the mean length of the lever. z is the mean vector of the class lever vectors (converter to CaM6) used as the z axis in **B** and **C**. **B:** Demonstration of how $r_c$ was calculated for each cryo-EM class using class A ($r_A$) and the distribution of end points from Fig. 4D **C:** Demonstration of how $x_c$ and $y_c$ were calculated for each cryo-EM class using class A ($x_A$ and $y_A$, respectively) and the distribution of end points from Fig. 4D. For the overall cantilever bending stiffness the displacement ($d_c$) was calculated using $r_c$ (**B**). To calculate the cantilever bending stiffness in each direction, tilt and slew, the displacement ($d_c$) was calculated using $x_c$ and $y_c$, respectively (**C**).

**Figure S4: Angles and distances between lever subdomain pairs**

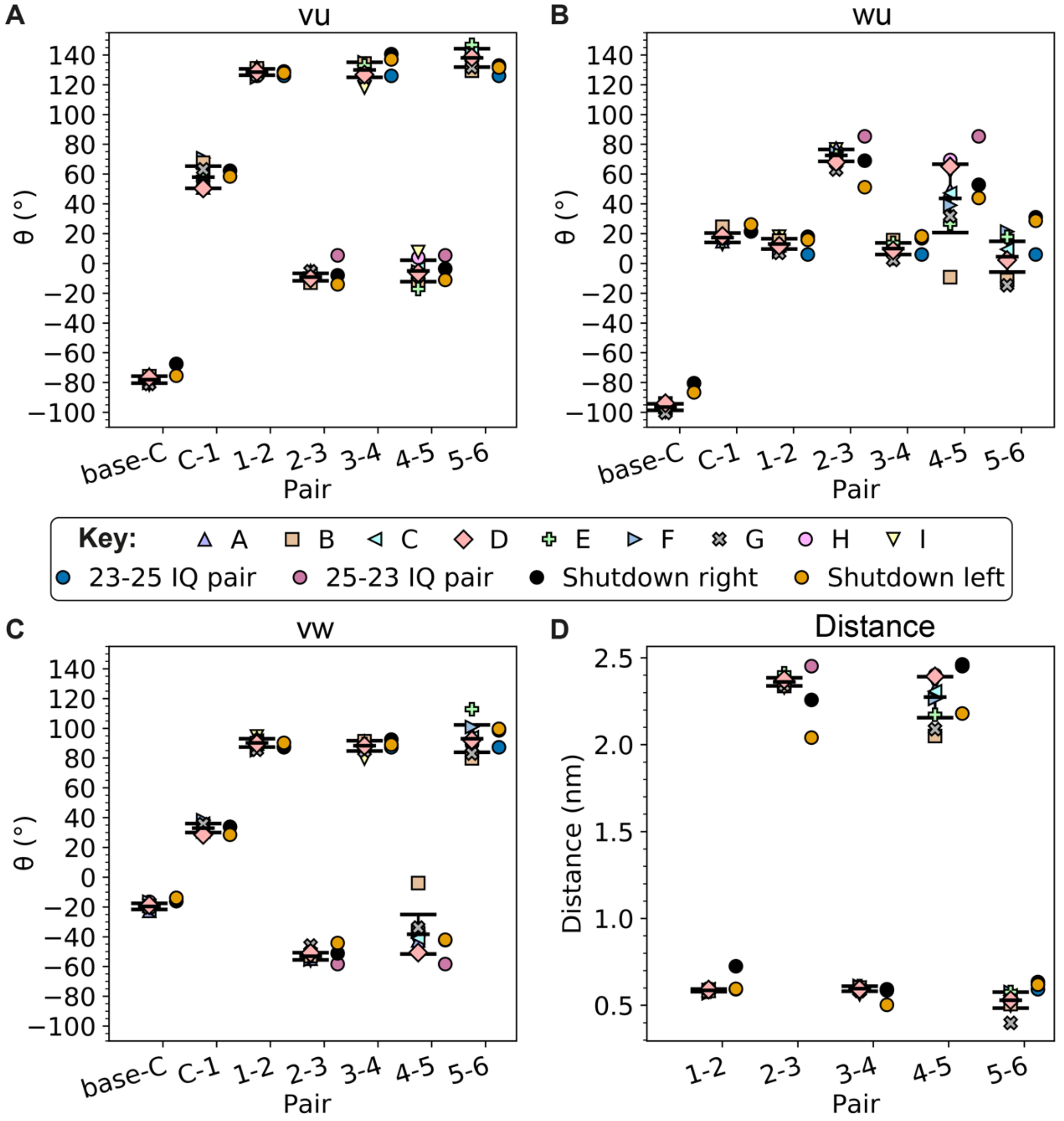


Classes A-H are named and colored as in Fig. 1A. Error bars show the mean and SD of the classes. Conformations of known structures of a 23-25 IQ pair bound to CaM, a 25-23 IQ pair bound to MLC1P and each subdomain in the shutdown state (PDB: 2IX7, 1N2D and 7YV9, respectively) [18,19] are displayed to the right of our data. Blue circles indicate values for the CaM pair bound to Myo5a IQ1-2 in the crystal structure (PDB: 2IX7, [19]). Pink circles indicate values for the MLC1P pair bound to Myosin-2p IQ2-3 in the crystal structure (PDB: 1N2D, [16]). Black circles indicate conformations of subdomain pairs in the structure of the shutdown state in the head furthest from the C-terminal region of the coiled-coil (right) (PDB: 7YV9 [18]). Orange circles indicate conformations of subdomain pairs in the structure of the shutdown state in the head closest to the C-terminal region of the coiled-coil (left) (PDB: 7YV9 [18]). **A:** The angle between lever subdomain vector pairs in their local vu plane (Fig. S5D). **B:** The angle between lever subdomain vector pairs in their local wu plane (Fig. S5E). **C:** The angle between lever subdomain vector pairs in their local vw plane (Fig. S5C). **D:** Distance between known interacting residues in the N- and C-lobes of consecutive CaM pairs (measured between the Cα of Ser17 and Asn111) [19]. Base = actin binding interface, C = converter, 1 = CaM1, 2 = CaM2, 3 = CaM4, 5 = CaM5, 6 = CaM6.

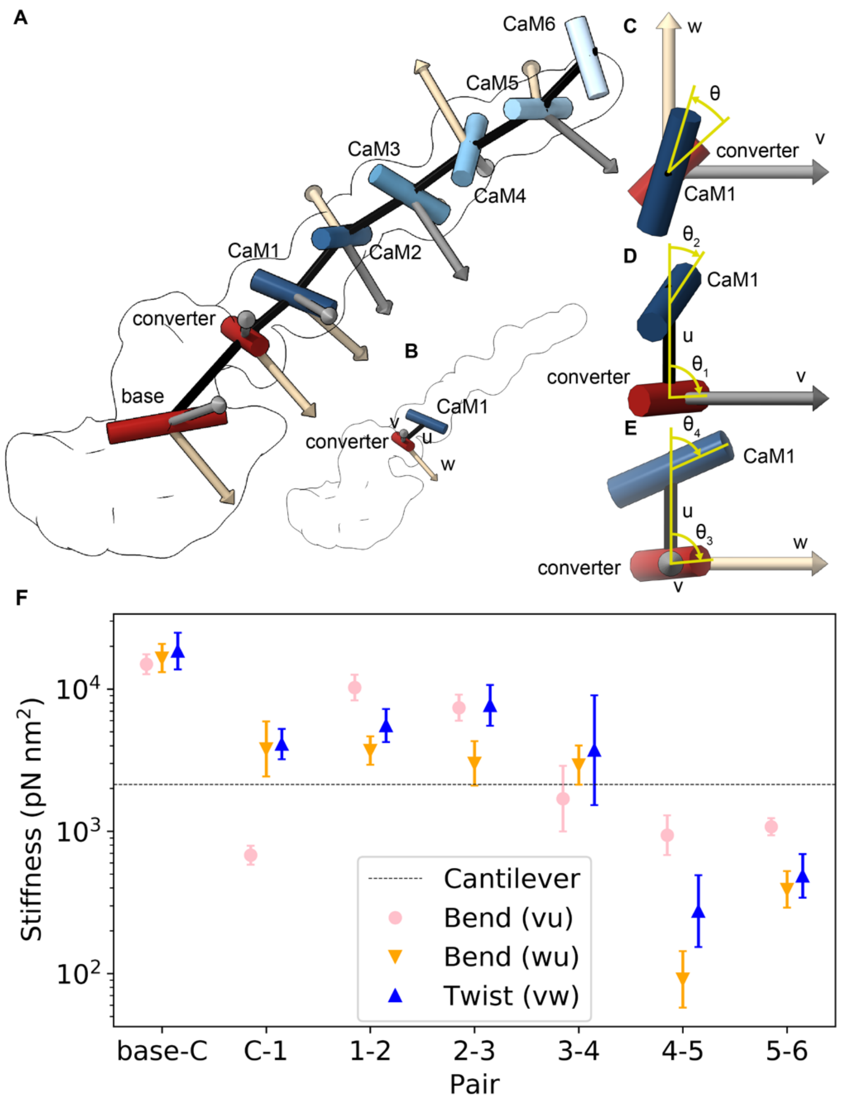


**Figure S5: Determining local subdomain directional bending and torsional stiffnesses from the cryo-EM data**

**A:** An example structure (class *d*) illustrates the subdomain vector pairs used in the calculation of local subdomain orientations (Fig. S4) and stiffnesses (**F**, Fig. 3B, Table S1), such as the base of the motor and the converter (base-C) and so on. Vectors representing key domains used for discretization (base, converter, CaMs 1-6) are shown as cylinders, and displayed within the gaussian filtered cryo-EM map. Local subdomain material axes (uvw) are shown as 3D arrows, where u is black, v is grey, and w is beige. **B:** As in **A** but focused on the subdomain comprising the converter and CaM1, with the local material axes labelled (uvw). **C:** An example of calculating the twist angle between a subdomain vector pair in the vw plane, using the vector representing the converter and the vector representing CaM1. $θ_{vw} = θ$. **D:** An example of calculating the angle between a subdomain vector pair in the vu plane, using the vector representing the converter and the vector representing CaM1. $θ_{vu} = θ_1$-$θ_2$. **E:** An example of calculating the angle between a subdomain vector pair in the wu plane, using the vector representing the converter and the vector representing CaM1. $θ_{wu} = θ_3$-$θ_4$. **F:** The calculated stiffnesses for subdomains as shown in **A**. The expected overall local bending stiffness for a beam with uniform bending along its length calculated from the cantilever bending stiffness is plotted as a black line (Cantilever). Bend (vu) is the bending stiffness in the vu plane. Bend (wu) is the bending stiffness in the wu plane. Twist (vu) is torsional stiffness in the vw plane. Base = actin binding interface, C = converter, 1 = CaM1, 2 = CaM2, 3 = CaM4, 5 = CaM5, 6 = CaM6. Error bars show the SD of the random error (see Materials and Methods for details) as a percentage of the reported stiffness.

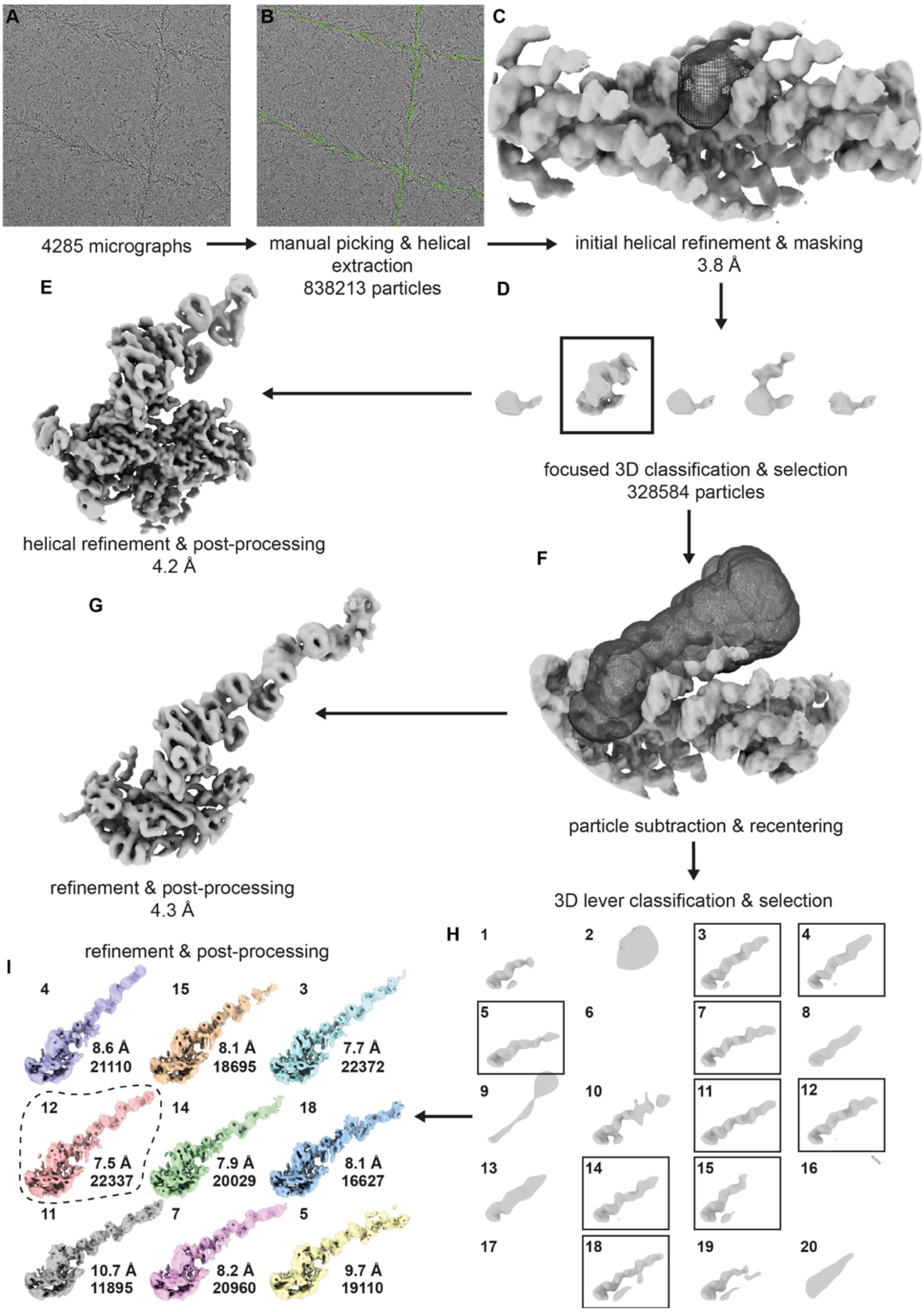


**Figure S6**: **Processing workflow**
Cryo-EM data processing pipeline for S1. **A**: Representative micrograph from 4285 micrographs. **B**: Helical picking of start end coordinates in Relion 3.1. **C**: Initial helical refinement in solid grey. Mask for 3D classification in black mesh. **D**: Resulting classes from focused 3D classification into 5 classes. Boxed class is the decorated F-actin class that was selected for further processing. **E**: 3D refined and post-processed map of motor + 2IQs. **F**: Cone subtraction mask in black mesh. Refined map of particles re-entered on subtraction mask coordinates in solid grey. **G**: 3D refined and post-processed map from particles where all density outside of the cone mask (**F**) has been subtracted. **H**: 3D classification into 20 classes within cone shaped lever mask. Boxed classes are the 9 classes with

density along the length of the lever selected for post-processing and analysis. **I**: Post-processed lever classes with number of particles in each class indicated below. All resolutions quoted are based on global resolution at 0.143 FSC (Fourier Shell Correlation). All resultant density maps (E, G & I) displayed were post-processed in DeepEMhancer [34].

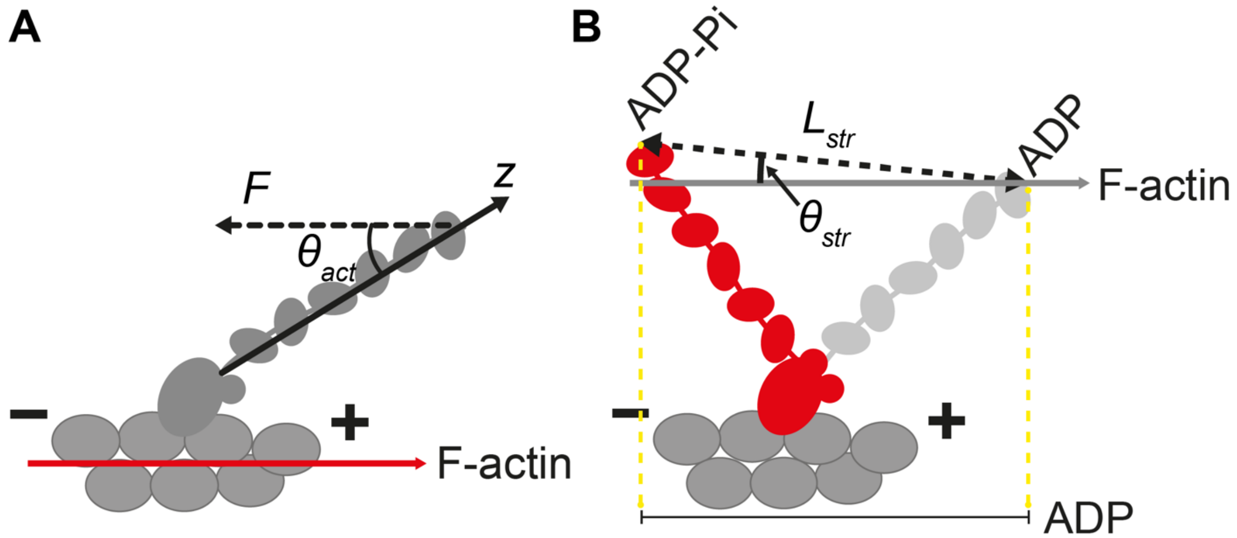


**Figure S7**: **Schematic diagrams to demonstrate how lever deflection and working stroke along F-actin are calculated.**
**A:** Schematic diagram to demonstrate how the applied torque was calculated ($F\sin\theta_{act}$). $F$ is the force applied, and $\theta_{act}$ is the angle between the mean lever position ($z$, as in Fig. 3C) and F-actin. **B:** Schematic diagram to demonstrate how each of the working strokes as a translation along F-actin were calculated. $L_{str}$ is the magnitude of an ADP-Pi to ADP lever end vector and $\theta_{str}$ is the angle between this vector and the F-actin axis. + indicates the plus end of F-actin, - indicates the minus end of F-actin.

**Table S1: Local subdomain stiffnesses (2.s.f), ± SD of random error (see Materials and Methods for details).**

| Subdomain | Combined bending stiffness (pN nm$^2$) | vu bending stiffness (pN nm$^2$) | wu bending stiffness (pN nm$^2$) | vw torsional stiffness (pN nm$^2$) |
|---|---|---|---|---|
| Base-converter | 16000 ± 3400 | 15000 ± 2600 | 16000 ±4200 | 18000 ± 6400 |
| converter-CaM1 | 2200 ± 1100 | 680 ± 110 | 3800 ± 2100 | 4100 ± 1100 |
| CaM1-2 | 7000 ± 1700 | 10000 ± 2400 | 3700 ± 950 | 5500 ± 1700 |
| CaM2-3 | 5200 ± 1500 | 7400 ± 1700 | 3000 ± 1300 | 7700 ± 3000 |
| CaM3-4 | 2300 ± 1100 | 1700 ± 1200 | 2900 ± 1100 | 3700 ± 5300 |
| CaM4-5 | 520 ± 200 | 940 ± 360 | 91 ± 53 | 270 ± 220 |
| CaM5-6 | 730 ± 150 | 1100 ± 160 | 390 ± 130 | 490 ± 210 |

**Table S2: Average subdomain lengths between cryo-EM 3D classes (2.s.f)**

| **Subdomain** | **Length (nm)** |
|---|---|
| Base-converter | 6.3 |
| converter-CaM1 | 2.9 |
| CaM1-2 | 3.5 |
| CaM2-3 | 3.7 |
| CaM3-4 | 3.5 |
| CaM4-5 | 3.8 |
| CaM5-6 | 3.3 |

### Table S3: Microscope parameters

| | |
|---|---|
| Microscope | Titan Krios I |
| Magnification | 75000 |
| Voltage (kV) | 300 |
| Electron dose per image ($e^-/Å^2$) | 62.66 |
| Exposure time (s) | 1.5 |
| Number of fractions | 59 |
| Defocus range (µm) | -1.8 to -3.6 (0.3 steps) |
| Pixel size (Å) | 1.065 |

**Table S4**: **MolProbity statistics for the pseudoatomic model of rigor Myo5a S1 bound to F-actin**

| **Atomic model statistics** | |
|---|---|
| MolProbity score | 0.73 |
| Clash score | 0.08 |
| Bad bonds (%) | 0.04 |
| Bad angles (%) | 0.55 |
| Poor rotamers (%) | 1.83 |
| Ramachandran favored (%) | 92.37 |
| Ramachandran outliers (%) | 0.11 |
| CaBLAM outliers (%) | 1.0 |
